# An unbiased comparison of immunoglobulin sequence aligners

**DOI:** 10.1101/2024.06.12.598680

**Authors:** Thomas Konstantinovsky, Ayelet Peres, Pazit Polak, Gur Yaari

**Author notes:** These authors contributed equally to this work (Thomas Konstantinovsky, Ayelet Peres).

## Abstract

Adaptive Immune Receptor Repertoire sequencing (AIRR-seq) is critical for our understanding of the adaptive immune system’s dynamics in health and disease. Reliable analysis of AIRR-seq data depends on accurate Immunoglobulin (Ig) sequence alignment. Various Ig sequence aligners exist, but there is no unified benchmarking standard representing the complexities of AIRR-seq data, obscuring objective comparisons of aligners across tasks. Here, we introduce GenAIRR, an efficient simulation framework for generating Ig sequences alongside their ground truths. GenAIRR realistically simulates the intricacies of V(D)J recombination, somatic hypermutation, and an array of sequence corruptions. We comprehensively assessed prominent Ig sequence aligners across various metrics, unveiling unique performance characteristics for each aligner. The GenAIRR-produced datasets, combined with the proposed rigorous evaluation criteria, establish a solid basis for unbiased benchmarking of immunogenetics computational tools. It sets up the ground for further improving the crucial task of Ig sequence alignment, ultimately enhancing our understanding of adaptive immunity.

## 1 Introduction

The adaptive immune system functionality relies upon a diverse and dynamic set of cell receptors. In lymphocytes, this diversity originates from the V(D)J recombination process [26], with B cells undergoing further diversification through affinity maturation; a process that includes clonal expansion [17], somatic hypermutation (SHM) [27], and affinity-dependent selection [45]. Advances in sequencing technologies, particularly adaptive immune receptor repertoire sequencing (AIRR-seq)[22], have profoundly enhanced our understanding of this repertoire, providing detailed insights into its dynamics and diversity in response to a wide spectrum of immunological challenges [42,41,19,9,2,15,37].

Analyzing AIRR-seq data requires an accurate alignment of immunoglobulin (Ig) sequences to their germline ancestors. This task poses significant computational challenges due to factors such as the vast array of known germline sequences [5], the stochastic nature of gene trimming during V(D)J recombination [39], alterations introduced by SHM [46], and ambiguities resulting from sequencing errors [44].

To address these challenges, two primary approaches are utilized for aligning Ig sequences: string distance metrics-based and Hidden Markov Models (HMM)-based. Distance-based methods [4, 48, 3], while computationally efficient, may encounter difficulties with complex sequence variations such as insertions, deletions, and mutations. In contrast, HMM-based methods [12, 25, 35] leverage probabilistic models to capture some of the stochastic nature of V(D)J recombination and sequence evolution. This approach provides a more detailed representation of the real diversity of Ig sequences and somatic evolutionary patterns. However, these methods can require more computational resources and rely on parameters inferred from empirical, potentially noisy, datasets.

All tools for Ig sequence alignment require a germline reference set that encompasses the known alleles expected to be included in AIRR-seq data. This germline reference set is used to establish the metrics necessary for the alignment. Current germline reference sets, such as those from IMGT [21] and OGRDB [20], suffer from either noise or incompleteness, further complicating alignment tasks. Thus, an adaptable germline reference set is essential for several reasons. First, numerous more recent studies have identified novel Ig alleles that are not present in standard reference databases (e.g., [14,23,36,24,13,28]). These newly identified alleles significantly contribute to immune repertoire diversity and play a crucial role in accurate alignment and analysis in personalized genomics [11, 10, 7, 35]. The importance of personalized genomics cannot be overstated, as individuals may have unique variations in their Ig genes, affecting immune responses and disease susceptibilities [6,30,8,34,47,1,18]. Furthermore, the ability to modify the reference to accommodate personalized genotypes ensures precise alignment and interpretation of Ig sequences. This adaptability also assists in identifying rare and low-frequency variants that may be critical to immune function but are often disregarded in standard reference-based alignments during immune repertoire analysis [32].

Understanding whether an Ig sequence is productive, or expressed, is vital for various aspects of immunological research. Since many factors contribute to the ability to express an Ig, only experimental validation can confirm the productivity status of a sequence. Nevertheless, several necessary conditions must be met for a sequence to be considered productive, which were identified originally by the International ImMunoGeneTics Information System (IMGT). The standardized framework to computationally infer the productivity of Ig sequences [33] includes ensuring a correct open reading frame; the absence of aberrations in the start codon, splicing sites, and regulatory elements; the absence of internal stop codons; and an in-frame junction region where the V, D, and J gene segments align properly. Despite these standardized criteria, variations in the assessment of sequence productivity can arise due to differences in algorithms and methodologies used by the different sequence aligners. Factors such as the handling of ambiguous gene segment boundaries, treatment of sequencing errors, and interpretation of junction regions can lead to discrepancies in sequence productivity classifications among aligners.

To address these challenges and accurately evaluate alignment tools, past benchmarks often used datasets derived from or simulated based on direct sequencing efforts [48, 40, 35]. These datasets inherently carry biases, like unequal allele representation and batch-effects prevalent in many cohorts. In addition, these benchmarks often overlooked critical aspects such as the ability of aligners to handle insertion or deletions (indels), accurately define the start and end positions of segments, and stratify performance across different levels of SHM. Further, the IGH/IGK/IGL genomic loci display high variability among individuals, posing challenges in creating representative reference sets. Such challenges underscore the need for a more comprehensive and unbiased benchmarking approach. The benefits of using objectively simulated data in such tasks is of paramount importance [38].

Building upon these insights, we propose a two-fold approach to address the challenges in benchmarking alignment tools effectively. First, we present a benchmarking setup that encompasses three critical metrics for evaluating sequence alignment tools: 1) Assessment of the tools’ accuracy in correctly identifying sequence allele calls. This precision is fundamental, as it forms the basis for understanding the alleles and genes at play, with significant downstream impacts on analyses such as genotype determination [29, 7], haplotype inference [31], cloning [44, 16], and SHM calls. 2) Segmentation: aiming to precisely identify the start and end of alleles within the sequence. Precision in this task is crucial because incorrect segmentation, such as premature or late trimming of the 3’ end of the V allele can lead to missed SHM events, influence the productivity assessment, or erroneous identification of non-real SHM events. 3) Productivity assessment of sequences. Downstream analysis pipelines commonly filter out what they consider to be non-productive sequences, hence correct assessment by the aligners has a high impact. Although evaluating productivity may seem straightforward, differences between aligners arise from variations in their algorithms and implementations. Second, we introduce GenAIRR, a robust simulation framework designed to generate Ig sequence datasets with established ground truths that enable accurate and comprehensive comparisons among aligners. GenAIRR incorporates realistic sequence corruptions and noise, filling gaps in existing simulation frameworks and providing a solid foundation for a robust benchmarking setup. See Supplementary Table 1 for an overview of existing simulation frameworks.

This manuscript aims not only to elevate the standards of aligner comparison, but also to establish a comprehensive framework for the ongoing evaluation of both existing and newly developed alignment methodologies. By doing so, it seeks to significantly improve the precision and reliability of AIRR-seq analysis. Such advancements will deepen our understanding of the adaptive immune system’s responses to pathogens and enhance our ability to leverage this knowledge in health and disease contexts.

## 2 Results

### 2.1 Creating a benchmarking setup using GenAIRR

To establish a robust benchmarking setup, a bias-free dataset is required. For this, we created GenAIRR, an AIRR-seq data simulator that simulates the full spectrum of V(D)J recombination events and introduces realistic sequence corruptions such as 5’ nucleotide trimming or addition, masking nucleotides with Ns, and introduction of indels. GenAIRR mitigates biases by enabling simulations without relying on empirical data distribution, opting instead for a uniform distribution to generate sequences (refer to Supplementary Table 1 for comparisons with other simulation tools). In addition, GenAIRR’s modular architecture allows users to tailor simulations to reflect specific experimental conditions. GenAIRR provides comprehensive ground truth data for each simulated sequence, including allele calls, segmentation positions, and productivity assessments, formatted in the AIRR community schema for annotated AIRR-seq data [43]. This facilitates straightforward comparisons with the output of commonly used alignment tools. GenAIRR is illustrated in Figure 1.

**Figure 1.**
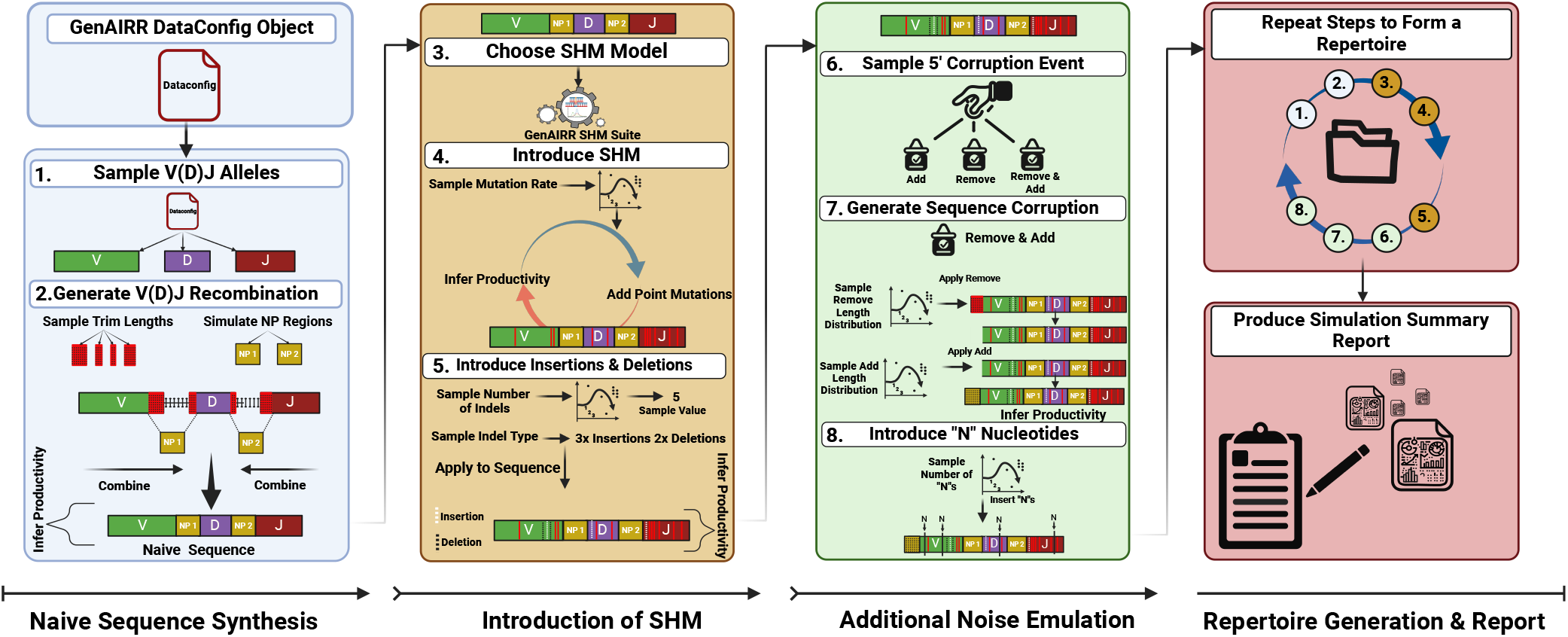
GenAIRR modular architecture to simulate Ig sequences. The first column (steps 1-2) describes simulation of a naive Ig sequence using the provided configuration file (Data-Config). The second column (steps 3-5) illustrates the introduction of alterations to the simulated sequence, such as SHM and indels. The third column (steps 6-7) illustrates the introduction of further experimental noise to the simulated sequence, including 5’ corruptions and N nucleotides. Finally, the fourth column illustrates that GenAIRR allows for repeating the sequence simulation to form a repertoire and generating a report summarizing its statistical properties.

Using GenAIRR, we created three datasets, each containing 6 million sequences, with a uniform distribution of alleles. The first dataset (DS1) consisted solely of productive sequences, devoid of corruptions, N masks, or indels, but did include varying mutation rates to mimic real AIRR-seq data. The second dataset (DS2) included mainly nonproductive sequences, resulting from the rearrangement process, introduction of mutations, or corruption events such as 5’ trimming or addition, N insertions, and indels (refer to Supplementary Table 3 for simulation details).

We used the GenAIRR report feature to check how alleles are distributed in these datasets. Although we aimed for an even distribution, we noticed a small difference in the usage of certain V and J alleles in DS1 (Fig. 2A and C) compared to DS2 (Supplementary Fig. 3A and C). This difference stems from constraints inherent in the generation process of productive sequences.

**Figure 2.**
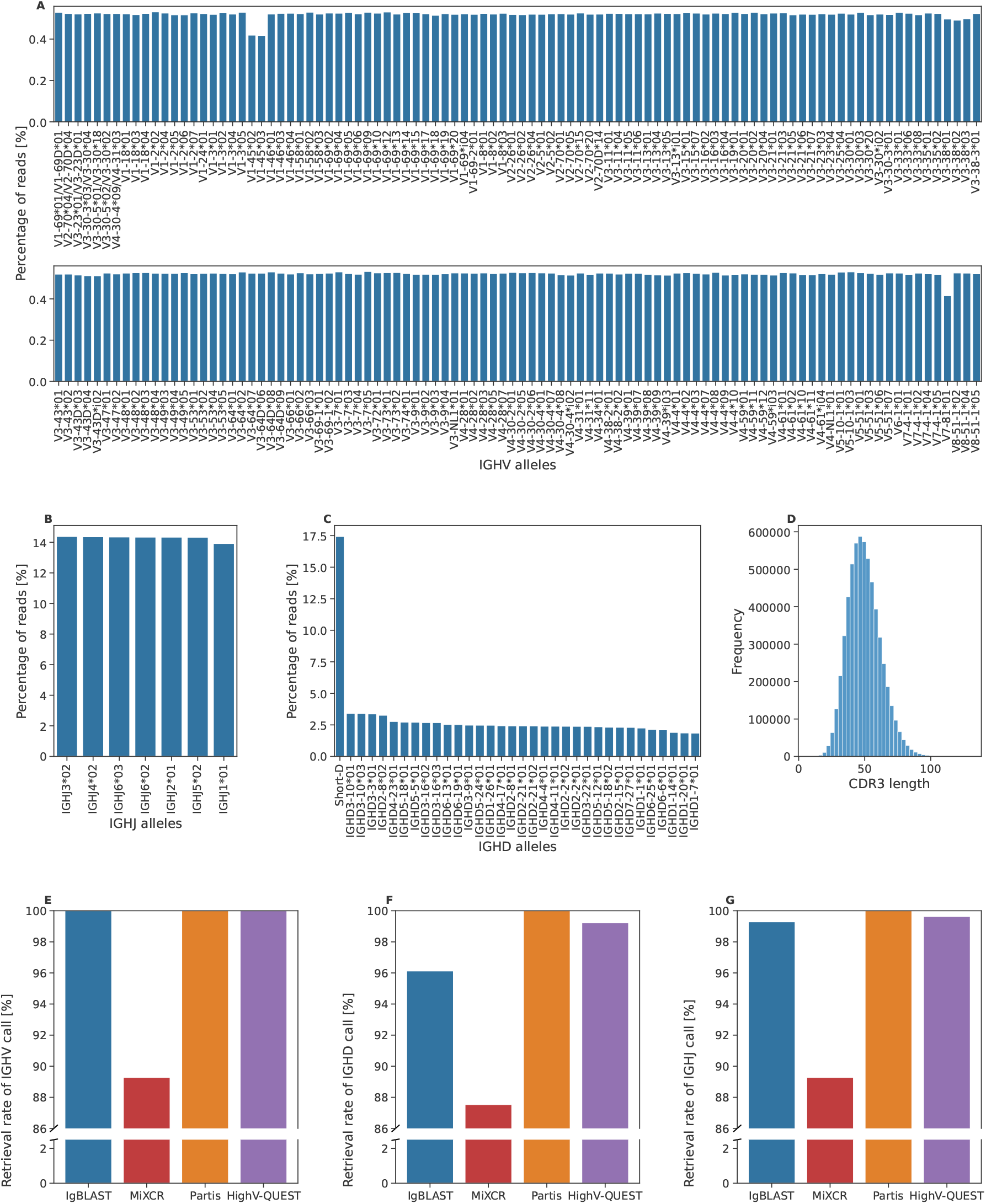
Overview of the simulated productive dataset. (A-C) Distribution of the V, D, and J allele usage in the productive dataset. Each column represents a different allele, and the y-axis indicates their relative usage percentage in the dataset. (D) CDR3 length distribution. The x-axis shows the CDR3 lengths, and the y-axis indicates their frequency. (E-G) Aligners allele assignment retrieval rate. Each column represents a different aligner, and the y-axis shows the percentage of sequences for which the aligner returned an allele assignment. The colors correspond to the different aligners: blue for IgBLAST, red for MiXCR, orange for Partis, and purple for HighV-QUEST.

In simulating D alleles, the protocol involved trimming both the 5’ and 3’ ends, sometimes resulting in very short sequences that pose alignment challenges due to their potential to match multiple alleles (see method section 4.2). In actual AIRR-seq data, it is impossible to ascertain the origin of these short D sequences. Hence, GenAIRR incorporates a feature that identifies sequences of five or fewer nucleotides as “Short-D” and conceals their origin. In both DS1 (Fig. 2B) and DS2 (Supplementary Fig. 3B), the allele usage across other D alleles was generally uniform. The simulated datasets exhibited a distribution of CDR3 lengths that resembled empirical data (Fig. 2D, Supplementary Fig. 3D).

### 2.2 Benchmarking immunoglobulin sequence aligners

In the presented benchmark evaluation, we first surveyed the existing alignment tools, and selected four widely used aligners, IgBLAST [48], MiXCR [3], HighV-QUEST [4], and Partis [35], based on their popularity, compatibility with the AIRR community schema for annotated AIRR-seq data [43], consistent support, and active development within the immunogenetics community (see Supplementary Table 2 for a summary). Importantly, these aligners were also chosen for their diverse alignment methodologies. IgBLAST employs BLAST for alignment, HighV-QUEST and MiXCR utilize multiple sequence alignment techniques, and Partis adopts an HMM approach.

Here, we utilized the newly published reference set by the AIRR community [5], which requires the aligners to enable the use of a custom reference set. HighV-QUEST is the only aligner of the four that is unable to accept a custom reference set, and thus in cases where the resulting allele assignment HighV-QUEST produced was not included in the AIRR-C reference set, we matched it with the closest allele in the set. Not all aligners return results for every sequence in the datasets. In DS1, IgBLAST, Partis, and HighV-QUEST provided assignments for all 6 million sequences at the V allele level, while MiXCR had a retrieval rate of approximately 89% (Fig. 2E). Furthermore, Partis consistently had a retrieval rate of 100% for both the D and the J alleles, closely followed by HighV-QUEST with ∼ 99%. The retrieval rate of IgBLAST for J alleles is roughly 99% and 96% for the D alleles. MiXCR showed a lower retrieval rate at 89% for J alleles and 87% for D alleles (Fig. 2F-G). The trend remained the same for the nonproductive sequences, with Partis returning the highest retrieval rate, trailed by IgBLAST, then IMGT, and lastly MiXCR (Supplementary Fig. 3E-G).

### 2.3 Precision of Aligners Varies with Segment Mutation Rates and Lengths

We evaluated the aligners’ ability to accurately assign V, D, and J allele calls for each sequence. We observed variations in performance across different mutation rates and sequence lengths (Fig. 3). IgBLAST demonstrated the highest overall accuracy in assigning V alleles. However, MiXCR excelled in sequences with a mutation rate of up to 10%, but rapidly declined at higher rates. Notably, HighV-QUEST exhibited underperformance at lower mutation rates but showed a relatively steady, good accuracy in highly mutated sequences (Fig. 3A).

**Figure 3.**
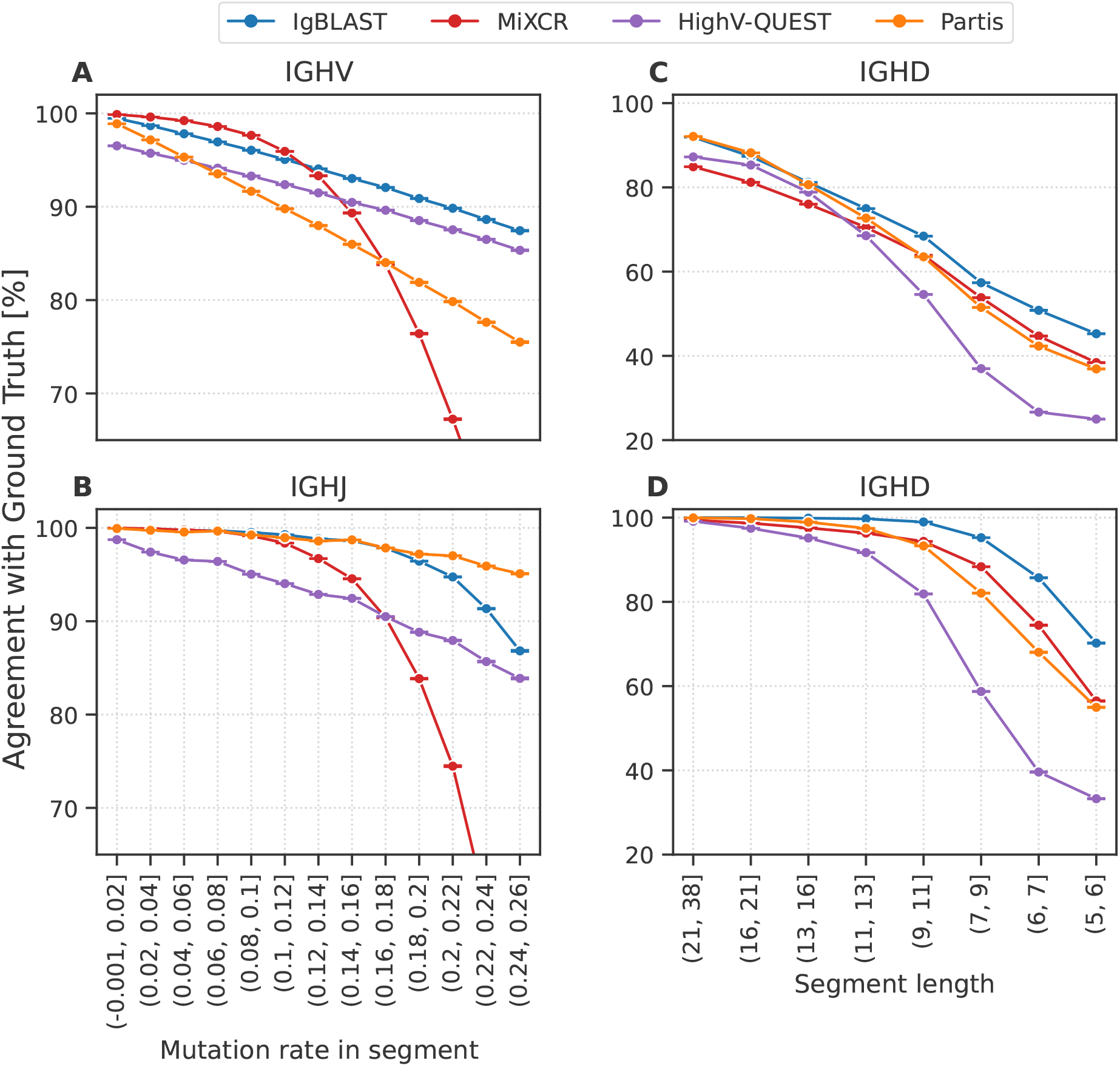
Aligners predictive performance of allele calls. (A+B) Agreement with ground truth for IGHV (A) and IGHJ (B) segment calls across varying mutation rates. The x-axis represents the mutation rates ranging from 0.01 to 0.25, and the y-axis shows the agreement percentage with the true allele call. The colors correspond to the aligners as indicated in the figure. (C) Agreement with ground truth for IGHD segment calls across varying segment lengths. The x-axis represents the varying D segment lengths from 38 to 6, and the y-axis shows the agreement percentage with the true allele call. (D) Same as C) for sequences with 0 mutations in their D segments.

Assigning the J allele was a relatively easier task due to the reduced variability among alleles. Partis consistently outperformed all aligners across various mutation rates, with IgBLAST closely following but showing a decline in sequences with over a 17% mutation rate. MiXCR maintained good accuracy up to a mutation rate of 11%, but, as with V alleles, its performance declined at higher mutation rates compared to other aligners (Fig. 3 B).

The evaluation of D assignment performance is shown as a function of D segment length rather than mutation rate. This is because mutation rates can vary significantly due to the short segment lengths, potentially masking the influence of both factors (Fig. 3C). For longer D sequences, Partis and IgBLAST exhibited slightly superior performance, with MiXCR lagging behind. For shorter D segments, IgBLAST remained the top performer, while HighV-QUEST exhibited the lowest agreement with ground truth. These findings underscore the nuanced performance dynamics of aligners depending on mutation rates and sequence lengths. To further assess the impact of segment length on accuracy, we isolated sequences that did not encounter any mutations within the IGHD segment (Fig. 3D). The accuracy of all aligners spiked across all segment lengths, with IgBLAST demonstrating accuracy above 90% for sequences longer than 9 nucleotides. However, as observed in Fig. 3C, accuracy declines in short D segments, emphasizing the influence of segment length on the ability to correctly assign the D allele.

### 2.4 Segmentation position accuracy differs among aligners

Besides accurately identifying alleles in AIRR-seq alignment, another crucial task is segmenting the sequence into distinct allele regions, which facilitates the correct attribution of SHM events. To evaluate the accuracy of segmentation by the aligners, we utilized both DS1 and DS2. We computed the Root Mean Square Error (RMSE) for each segment’s 5’ and 3’ positions across all aligners. To minimize bias, we only considered perfect matches between allele calls and the ground truth, prioritizing the first matching call in cases of multiple allele calls and excluding V 5’ and J 3’ segmentations for this analysis.

In DS1, the RMSE results for the V 3’ position were generally similar across the aligners, except MiXCR, which exhibited a higher value (Fig. 4A). The accuracy in segmenting the D 5’ position was consistent across all aligners. However, for the D 3’ position, Partis showed a slightly higher RMSE value compared to the others. Regarding the J 5’ position, HighV-QUEST recorded the highest RMSE, while Partis demonstrated the lowest value. In DS2, the aligners maintained relatively similar RMSE and ranking values, as observed in DS1 (Fig. 4B). These findings underscore the significant disparities in segmentation accuracy across aligners and their performance across different dataset contexts.

**Figure 4.**
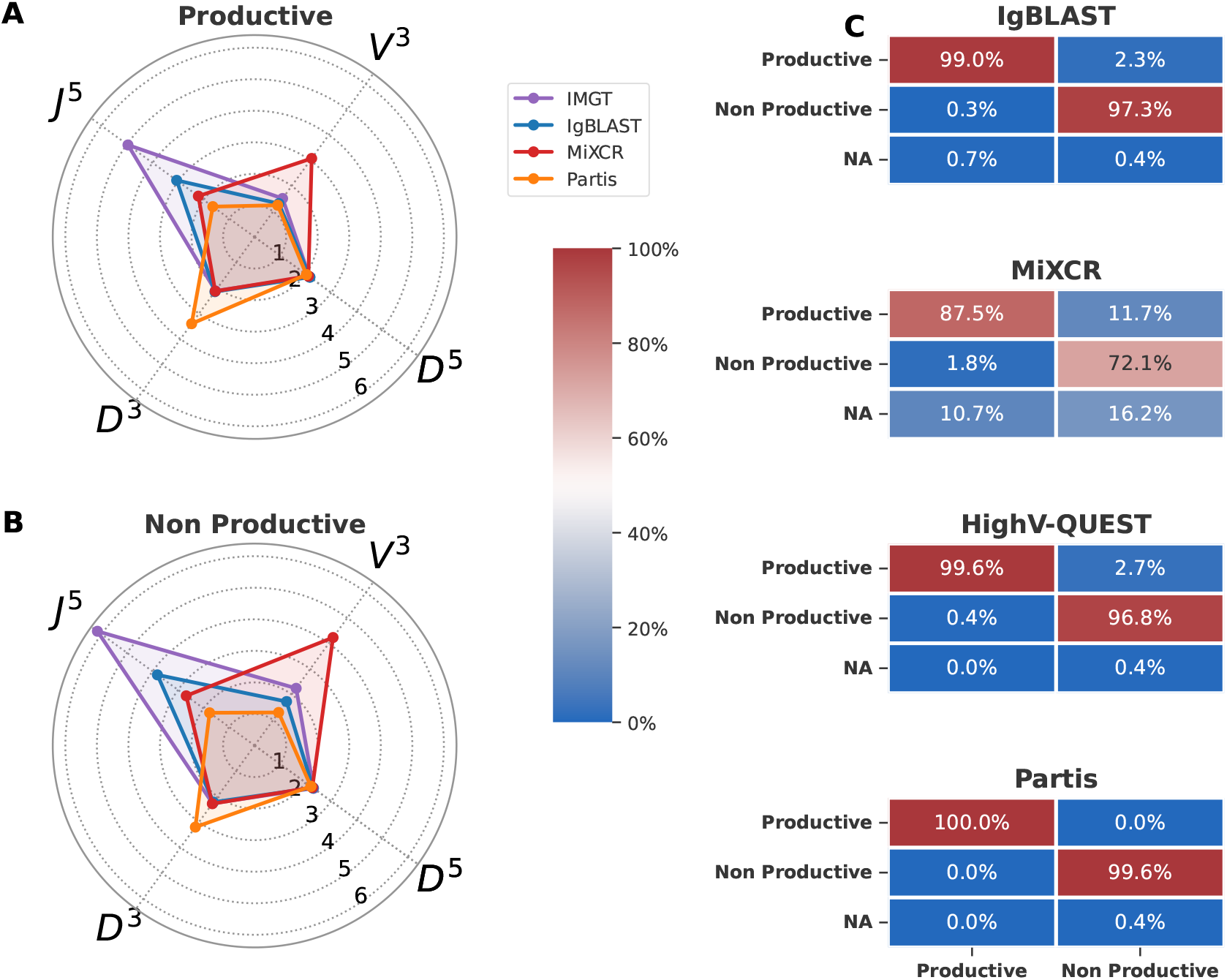
Aligners segmentation and productivity assessment. (A+B) Radar charts illustration of the RMSE (Root Mean Square Error) values for segmentation accuracy across gene segment start and end positions (V3, D3, D5, J3) in productive (A) and non-productive (B) sequences. (C) The confusion matrices provide an accuracy comparison of productive assessments. In each matrix, columns represent the 4M sequence subset of the simulated dataset, while rows represent the aligners’ productive assessment categorized as Productive, Non Productive, or NA (indicating no assessment for the sequence). The value in each cell indicates the percentage of agreement with the ground truth. The color scale reflects the level of accuracy.

### 2.5 Aligners Accuracy in Determining Sequence Productivity

Sequence productivity is important for immune repertoire analysis tools, affecting their precision in tasks such as genotype, haplotype, clonality, and diversity assessments. To assess the accuracy of the aligners in evaluating sequence productivity, we initially established the ground truth using GenAIRR. GenAIRR determines sequence productivity based on the following criteria: absence of a stop codon, presence of conserved AA at the start and end of the junction, and alignment that is in-frame (Supplementary Figure 2). To assess aligner accuracy in determining productivity, we used dataset DS3, which contains four million randomly sampled productive sequence from DS1, and a matching number of randomly sampled nonproductive sequence from DS2.

For productive sequences, Partis demonstrated perfect accuracy, whereas IgBLAST showed a low error rate with a slightly higher rate of missing sequences, marginally outperforming HighV-QUEST, which had a higher error rate but no missing sequences. Lastly, MiXCR exhibited a higher error rate and the most substantial rate of missing sequences (Fig. 4C, left column). The pattern was comparable for nonproductive sequences. Partis showed the highest accuracy, with a small proportion of sequences missing. IgBLAST performed slightly better than HighV-QUEST in terms of error rates. MiXCR exhibited a high error rate and the largest proportion of missing sequences (Fig. 4C right column).

## 3 Discussion

The current study represents an unbiased evaluation of Ig sequence aligners, facilitated by the GenAIRR simulation framework. This comprehensive assessment across multiple metrics has outlined distinct strengths and weaknesses inherent in the widely used aligners IgBLAST, MiXCR, HighV-QUEST, and Partis. The datasets generated by GenAIRR provide a robust, unbiased platform for benchmarking these aligners in a spectrum of challenging immunogenetic variables, mimicking real-world complexities. This benchmarking analysis explores how noise levels, such as mutation rates and corruption events, affect aligner agreement with the ground truth. The aligners’ comparisons (Table 1) revealed varying performances. MiXCR emerged as the most efficient aligner in terms of runtime and was the top performer for V assignment accuracy at low mutation rates (<10%), but decreased dramatically at higher mutation rates. Moreover, it showed lower accuracy for D assignment across all segment lengths. IgBLAST, although >10 times slower than MiXCR, outperformed the other aligners in almost all allele assignment accuracies. One exception is Partis, which demonstrated excellent performance in J assignment accuracy across both low and high mutation rates. The accuracy metric used here for allele assignment is relatively lenient, requiring a single assignment per sequence to match the ground truth. This leniency can disadvantage aligners that, by default, return only one assignment, such as Partis, as they have a lower probability of matching the ground truth. Note that erroneous assignments can in many cases be rectified by inferring a personal genotype.

**Table 1:** Comparison of different IG sequence aligners focusing on criteria such as custom reference acceptance, operational efficiency, and accuracy across mutation rates (MR). Accuracy metrics are presented for productive (left) and non-productive (right) sequences, with the best results highlighted in **bold**. RMSE (Root Mean Square Error) quantifies the precision of segment start and end position predictions; lower values indicate higher precision.

| Criteria | IgBLAST |  | MiXCR |  | HighV-QUEST |  | partis |  |
| --- | --- | --- | --- | --- | --- | --- | --- | --- |
|  | Prod. | Non | Prod. | Non | Prod. | Non | Prod. | Non |
| **Accepts Custom Reference** | **✓** |  | **✓** |  | **✗** |  | **✓** |  |
| **6M Seqs Runtime (Minutes)** | 2200 |  | **200** |  | — |  | 5400 |  |
| **V Accuracy (MR < 10%)** | 97.79 | 97.57 | **98.99** | **97.78** | 94.21 | 94.61 | 95.31 | 93.43 |
| **V Accuracy (MR > 10%)** | **90.78** | **89.99** | 75.09 | 66.92 | 88.45 | 88.18 | 81.77 | 80.79 |
| **J Accuracy (MR < 10%)** | **99.74** | **99.61** | 99.69 | 99.34 | 96.83 | 95.14 | 99.62 | 99.24 |
| **J Accuracy (MR > 10%)** | 93.69 | 92.49 | 70.86 | 68.85 | 88.48 | 86.58 | **97.0** | **96.86** |
| **D Accuracy (Length < 10)** | **51.14** | **49.42** | 45.63 | 42.44 | 29.53 | 26.73 | 43.57 | 42.47 |
| **D Accuracy (Length > 10)** | **85.3** | **84.16** | 79.26 | 74.75 | 80.36 | 75.52 | 84.69 | 83.81 |
| **V 3' RMSE** | 1.31 | 1.72 | 3.08 | 4.23 | 1.51 | 2.24 | **1.24** | **1.29** |
| **D 5' RMSE** | 2.17 | 2.29 | 2.11 | 2.26 | 2.15 | 2.31 | **2.03** | **2.2** |
| **D 3' RMSE** | 2.14 | **2.22** | 2.12 | 2.28 | **2.11** | 2.27 | 3.4 | 3.2 |
| **J 5' RMSE** | 3.06 | 3.81 | 2.2 | 2.68 | 4.96 | 6.16 | **1.64** | **1.76** |
| **Productivity Accuracy** | 99 | 97.3 | 87.5 | 72.1 | 99.6 | 96.8 | **100** | **99.6** |

Properly segmenting a sequence can influence factors such as attributing SHM events to the correct segments and assessing sequence productivity. Partis showed the lowest RMSE for position predictions in most categories, and the highest accuracy in the productivity assessment. Error rates in productivity can be attributed not only to segmentation errors, which can lead to missed stop codons, but also to misclassifying sequences that lack conserved residues in the junction as productive.

GenAIRR’s modularity is a standout feature, allowing the simulation to start from a naive sequence and progressively introduce noise (Table 2). This modular design facilitates easy adaptation of the code to different noise models and addition of optional stages, such as simulating epigenetic modifications, incorporating complex recombination patterns, or adding sophisticated SHM models. GenAIRR is released as an open source code, to allow community contributions to the simulation framework that may enhance its versatility and utility beyond the application presented here. Here, we used GenAIRR to simulate human Ig sequences, but other species can present different challenges. For example, macaque monkeys have many more known V(D)J alleles [49] compared to human, and as such can affect aligner performances in a non-trivial manner. In conclusion, our study not only presents a robust benchmarking setup for refining existing alignment tools and encouraging the development of new ones, but also underscores the immense potential of GenAIRR’s modular and adaptable framework. By integrating advanced functionalities and addressing key challenges in simulating Ig sequences, we pave the way for more accurate and reliable bioinformatics tools in immunogenetics. This collaborative effort, coupled with standardized benchmarking criteria using simulated sequences with known ground truth, propels the field towards optimized algorithms and deeper insights into immune system analysis, ultimately benefiting healthcare and disease research.

**Table 2:**
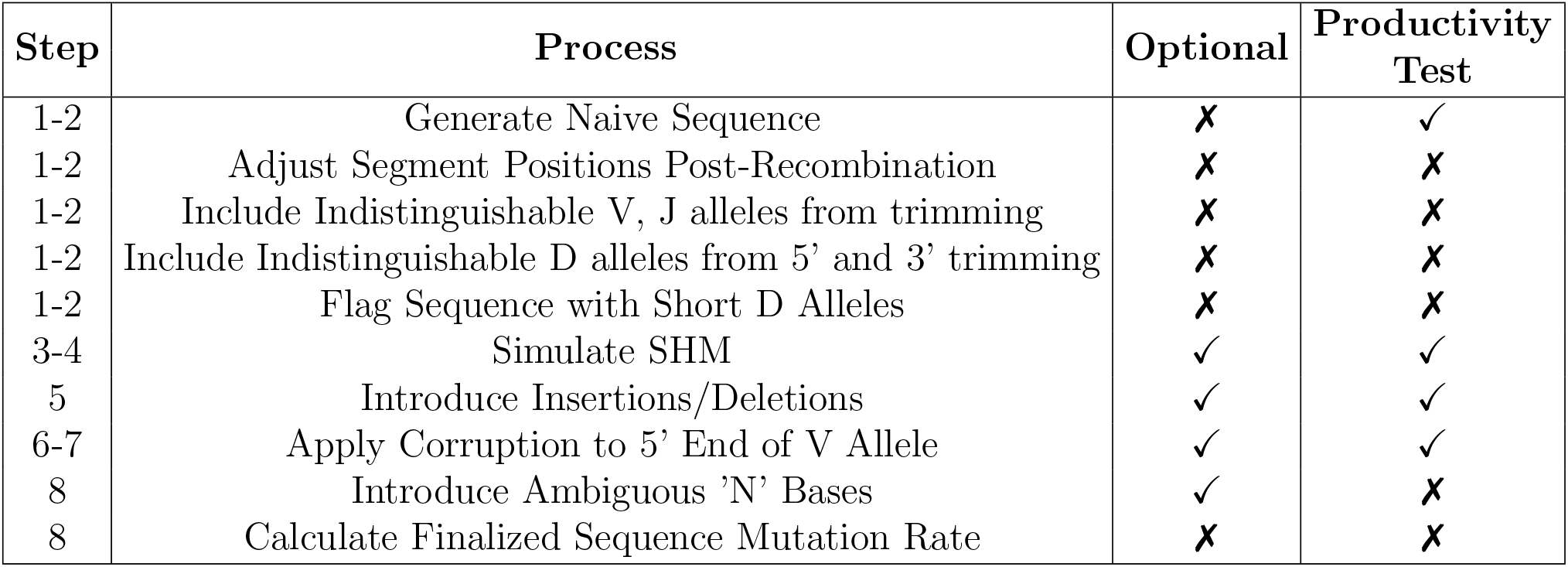
Illustration of the main GenAIRR sequence simulation workflow. Each numbered step corresponds to a stage in Figure 1. Steps marked as ‘Optional’ can be adjusted or omitted based on specific simulation needs, and steps marked under ‘Productivity Test’ indicate whether a productivity assessment is performed at this step.

## 4 Methods (online)

### 4.1 Simulation Workflow of GenAIRR

The GenAIRR simulation framework consists of a series of customizable modular events designed to generate Ig sequences that reflect the complexity and variability found in natural immune responses. Each event is controlled by a set of adjustable parameters, providing a high degree of flexibility and coverage.

The simulation begins with a stochastic selection of variable (V), diversity (D), and joining (J) alleles from a representative pool. In the context of the current manuscript, the aim was to generate datasets with a uniform representation of all alleles. Once selected, the allele sequences undergo specific trimming processes to enhance their structural complexity. Additionally, the nontemplated, palindromic (NP) regions are integrated using Markov chains derived from empirical data, which model the probabilistic arrangement of nucleotides (see Supplementary Section 1). Next, the simulation incorporates controlled mutation rates, models of SHM, and indels to enrich sequence diversity. GenAIRR also provides the option to introduce ambiguous ‘N’ bases deliberately, to test the robustness of sequence alignment algorithms. Additionally, sequencing artifacts are simulated, particularly at the 5’ ends of V alleles to mimic shorter sequences and 5’ untranslated region residuals. These features are introduced through tunable parameters that control the type, probability, and extent of errors. A detailed overview of the simulation steps is provided in Table 2, and an example of the tunable parameters used in this manuscript is provided in Supplementary Table 3.

### 4.2 Ground Truths Produced by GenAIRR

GenAIRR generates detailed ground truth data for each Ig sequence, which is essential for evaluating the accuracy of sequence alignment tools. These data include information on the V, D, and J allele calls, and their respective segment start and end positions. In addition, the ground truth includes documentation of the mutations, trimmings, indels, productivity, and corruption events (see Supplementary Table 4 for ground truth example).

During the simulation of Ig sequences, GenAIRR incorporates several strategies to address challenges such as the potential ambiguities arising from trimming or corruption at the allele ends. One significant challenge is the short length of D alleles, caused by their substantial trimming at both the 5’ and 3’ ends during recombination. This often results in sequences retaining only a minimal number of bases, making it impossible to distinguish these short sequences from multiple D alleles. GenAIRR manages this by employing a hyperparameter threshold to determine the minimal length for a D allele to be considered distinguishable post-recombination. Alleles shorter than this threshold are labeled as “Short-D”. The chosen threshold of five bases reflects a balance between computational efficiency and biological realism, covering approximately 85% of D alleles, as seen in Supplementary Figure 1. A map object post-trimming identifies which alleles become indistinguishable under specific scenarios, and adjustments to the ground truth are made. This includes corrections to the start and end positions of each allele following recombination and the simulated insertion of NP regions to verify that the trimmed section was not partially reconstructed by the NP regions. Such careful adjustments maintain the integrity of the ground truth, enabling more precise and reliable simulation outcomes (see Supplementary Section 2.1).

In addition to addressing these challenges, GenAIRR assesses the productivity of the simulated sequence. The criteria for a productive sequence are the absence of stop codons, the sequence to be in the correct open reading frame, and the presence of two conserved amino acids (AA) before and after the Complementary Determining Region 3 (CDR3). The region encompassing these conserved AAs and CDR3 is termed the junction. Sequence productivity is assessed four times during the GenAIRR sequence simulation (refer to Table 2, and Supplementary Fig. 2).

### 4.3 Benchmarking setup

For creating our benchmarking setup, we have generated three datasets, created four comparison matrices, and selected suitable alignment tools for comparison.

#### 4.3.1 Data Preparation

Using GenAIRR, two datasets, each containing 6 million sequences, were simulated for this study. These datasets were generated using the AIRR-C reference set [5] and clustered using the Allele Similarity Cluster method [32] to remove identical sequences. Within this reference set, there are 192 V alleles, 33 D alleles, and 7 J alleles. The selection of alleles for the simulated sequences followed a uniform distribution. This approach aimed to achieve an equal representation of all alleles in the reference and to eliminate any potential biases.

The first dataset, DS1, comprised purely productive sequences, reflecting common AIRR-seq data, enabling us to evaluate alignment tools under typical conditions. Supplementary Table 3 shows the parameters used to generate this dataset. The second dataset, DS2, primarily consisted of non-productive sequences resulting from recombination, SHM events, or noise introduction (Supplementary Table 3). This allowed us to test the alignment tools under extreme conditions and assess their performance. The third dataset, DS3, was used to assess productivity calls in a combination of data from DS1 and DS2. In summary:

- **DS1**: Includes only productive sequences without introduced corruptions, representing optimal alignment scenarios.
- **DS2**: Contains sequences with intentional corruptions like ambiguous base maskings (‘N’s), 5’ corruption events (removal or addition of nucleotides), and indels. This dataset comprises mostly non-productive sequences, with a minor fraction of productive sequences.
- **DS3**: A combined dataset comprising sampled sequences from both DS1 and DS2. Specifically, four million productive sequences were randomly sampled from DS1, along with four million non-productive sequences from DS2.

### 4.4 Benchmarking Immunoglobulin Sequence Alignment tools

In this study, we surveyed the existing alignment tools (Supplementary Table 2), and selected four popular tools for comparison. These tools were selected not only because of their popularity, but also because they are consistently maintained and developed in the immunogenetics community, and also comply with the AIRR community schema for annotated AIRR-seq data [43]. The leading aligners that were evaluated here are IgBLAST[48], MiXCR[3], HighV-QUEST[4] by IMGT, and Partis[35].

#### 4.4.1 Benchmarking Criteria

To benchmark the performance of the aligners, we have used four metrics:

- **Alignment Accuracy**: The effectiveness with which each aligner identified V, D, and J alleles under varying conditions of mutation rates and sequence corruptions was evaluated. Accuracy was quantified using an “Agreement” metric, defined for each sequence as 1 if the intersection of ground truth alleles (G) and alleles predicted by the aligner (P) is not empty *G*∩ *P* ≠ ∅, and 0 otherwise. This metric directly measures the ability of an aligner to correctly identify alleles used to generate the sequence.
- **Segmentation Accuracy**: We evaluated aligners’ precision in segmenting gene regions, focusing on the accuracy of segment start and end positions amidst mutations and sequence corruptions. This assessment is restricted to sequences for which there was a perfect match between the aligners’ allele calls for V, D, and J and the ground truth. In cases where the aligner made multiple assignments for any of V, D, or J alleles, the sequence was included in the comparison only if the first assignment matched the ground truth. This approach was adopted because aligners’ segmentation values are associated with the first assignment; hence, including sequences with erroneous values from subsequent assignments would skew the analysis. An adjustment was made for Partis due to intentional N-padding at the beginning of sequences. The results were modified to align Partis’ segmentation positions accurately with the ground truth by subtracting the added N-padding. To quantify segmentation errors, we utilized the Root Mean Square Error (RMSE) metric, defined as:

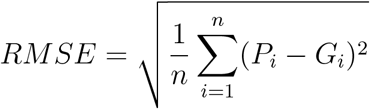

where *n* is the number of sequences, *P*_*i*_ is the predicted segment position by an aligner, and *G*_*i*_ is the actual ground truth position.
- **Productive Sequence Detection Accuracy**: This evaluation involved the calculation of the proportion of true positives and true negatives, which were then utilized to determine the overall classification accuracy. The results are presented using confusion matrices.
- **Runtime Efficiency and Resource Utilization**: The time needed to process 6 million sequences was measured for each aligner to compare computational efficiency. This evaluation was conducted on our cluster with specifications including an Intel(R) Xeon(R) Gold 6130 CPU @ 2.10GHz, 64 CPU cores, and 376 GB of RAM. The aligners were run on a single thread without parallelization, to align 60,000 sequences. The processing time for this subset was then multiplied by 100 to estimate the runtime for 6 million sequences.

## References

[1] Yuval Avnir, Aimee S Tallarico, Quan Zhu, Andrew S Bennett, Gene Connelly, Jared Sheehan, Jianhua Sui, Amr Fahmy, Chiung-yu Huang, Greg Cadwell, et al. Molecular signatures of hemagglutinin stem-directed heterosubtypic human neutralizing antibodies against influenza a viruses. PLoS pathogens, 10(5):e1004103, 2014.

[2] RJM Bashford-Rogers, Laura Bergamaschi, EF McKinney, DC Pombal, Federica Mescia, JC Lee, DC Thomas, SM Flint, P Kellam, DRW Jayne, et al. Analysis of the b cell receptor repertoire in six immune-mediated diseases. Nature, 574(7776):122–126, 2019.

[3] Dmitriy A Bolotin, Stanislav Poslavsky, Igor Mitrophanov, Mikhail Shugay, Ilgar Z Mamedov, Ekaterina V Putintseva, and Dmitriy M Chudakov. Mixcr: software for comprehensive adaptive immunity profiling. Nature Methods, 12(5):380–381, April 2015.

[4] X. Brochet, M.-P. Lefranc, and V. Giudicelli. Imgt/v-quest: the highly customized and integrated system for ig and tr standardized v-j and v-d-j sequence analysis. Nucleic Acids Research, 36(Web Server):W503–W508, May 2008.

[5] Andrew M Collins, Mats Ohlin, Martin Corcoran, James M Heather, Duncan Ralph, Mansun Law, Jesus Martínez-Barnetche, Jian Ye, Eve Richardson, William S Gibson, et al. Airr-c ig reference sets: curated sets of immunoglobulin heavy and light chain germline genes. Frontiers in Immunology, 14:1330153, 2024.

[6] Andrew M Collins, Gur Yaari, Adrian J Shepherd, William Lees, and Corey T Watson. Germline immunoglobulin genes: disease susceptibility genes hidden in plain sight? Current Opinion in Systems Biology, 24:100–108, 2020.

[7] Martin M Corcoran, Ganesh E Phad, Néstor Vázquez Bernat, Christiane Stahl-Hennig, Noriyuki Sumida, Mats AA Persson, Marcel Martin, and Gunilla B Karlsson Hedestam. Production of individualized v gene databases reveals high levels of immunoglobulin genetic diversity. Nature communications, 7(1):13642, 2016.

[8] Allan C Decamp, Martin M Corcoran, William J Fulp, Jordan R Willis, Christopher A Cottrell, Daniel LV Bader, Oleksandr Kalyuzhniy, David J Leggat, Kristen W Cohen, Ollivier Hyrien, et al. Human immunoglobulin gene allelic variation impacts germline-targeting vaccine priming. npj Vaccines, 9(1):58, 2024.

[9] Ali H Ellebedy, Katherine JL Jackson, Haydn T Kissick, Helder I Nakaya, Carl W Davis, Krishna M Roskin, Anita K McElroy, Christine M Oshansky, Rivka Elbein, Shine Thomas, et al. Defining antigen-specific plasmablast and memory b cell subsets in human blood after viral infection or vaccination. Nature immunology, 17(10):1226–1234, 2016.

[10] Daniel Gadala-Maria, Moriah Gidoni, Susanna Marquez, Jason A. Vander Heiden, Justin T. Kos, Corey T. Watson, Kevin C. O’Connor, Gur Yaari, and Steven H. Kleinstein. Identification of subject-specific immunoglobulin alleles from expressed repertoire sequencing data. Frontiers in Immunology, 10:129, 2019.

[11] Daniel Gadala-Maria, Gur Yaari, Mohamed Uduman, and Steven H. Kleinstein. Automated analysis of high-throughput b-cell sequencing data reveals a high frequency of novel immunoglobulin v gene segment alleles. Proceedings of the National Academy of Sciences, 112(8):E862–E870, 2015.

[12] Bruno A. Gäeta, Harald R. Malming, Katherine J.L. Jackson, Michael E. Bain, Patrick Wilson, and Andrew M. Collins. ihmmune-align: hidden markov model-based alignment and identification of germline genes in rearranged immunoglobulin gene sequences. Bioinformatics, 23(13):1580–1587, April 2007.

[13] William S Gibson, Oscar L Rodriguez, Kaitlyn Shields, Catherine A Silver, Abdullah Dorgham, Matthew Emery, Gintaras Deikus, Robert Sebra, Evan E Eichler, Ali Bashir, et al. Characterization of the immunoglobulin lambda chain locus from diverse populations reveals extensive genetic variation. Genes & Immunity, 24(1):21–31, 2023.

[14] Moriah Gidoni, Omri Snir, Ayelet Peres, Pazit Polak, Ida Lindeman, Ivana Mikocziova, Vikas Kumar Sarna, Knut EA Lundin, Christopher Clouser, Francois Vigneault, et al. Mosaic deletion patterns of the human antibody heavy chain gene locus shown by bayesian haplotyping. Nature communications, 10(1):1–14, 2019.

[15] Miri Gordin, Hagit Philip, Alona Zilberberg, Moriah Gidoni, Raanan Margalit, Christopher Clouser, Kristofor Adams, Francois Vigneault, Irun R Cohen, Gur Yaari, et al. Breast cancer is marked by specific, public t-cell receptor cdr3 regions shared by mice and humans. PLoS Computational Biology, 17(1):e1008486, 2021.

[16] Namita T Gupta, Jason A Vander Heiden, Mohamed Uduman, Daniel Gadala-Maria, Gur Yaari, and Steven H Kleinstein. Change-o: a toolkit for analyzing large-scale b cell immunoglobulin repertoire sequencing data. Bioinformatics, 31(20):3356–3358, 2015.

[17] Philip D Hodgkin, William R Heath, and Alan G Baxter. The clonal selection theory: 50 years since the revolution. Nature immunology, 8(10):1019–1026, 2007.

[18] Todd A Johnson, Yoichi Mashimo, Jer-Yuarn Wu, Dankyu Yoon, Akira Hata, Michiaki Kubo, Atsushi Takahashi, Tatsuhiko Tsunoda, Kouichi Ozaki, Toshihiro Tanaka, et al. Association of an ighv3-66 gene variant with kawasaki disease. Journal of human genetics, 66(5):475–489, 2021.

[19] Uri Laserson, Francois Vigneault, Daniel Gadala-Maria, Gur Yaari, Mohamed Uduman, Jason A Vander Heiden, William Kelton, Sang Taek Jung, Yi Liu, Jonathan Laserson, et al. High-resolution antibody dynamics of vaccine-induced immune responses. Proceedings of the National Academy of Sciences, 111(13):4928–4933, 2014.

[20] William Lees, Christian E Busse, Martin Corcoran, Mats Ohlin, Cathrine Scheepers, Frederick A Matsen IV, Gur Yaari, Corey T Watson, AIRR Community, Andrew Collins, et al. Ogrdb: a reference database of inferred immune receptor genes. Nucleic acids research, 48(D1):D964–D970, 2020.

[21] Marie-Paule Lefranc, Veronique Giudicelli, Chantal Ginestoux, Joumana Jabado-Michaloud, Geraldine Folch, Fatena Bellahcene, Yan Wu, Elodie Gemrot, Xavier Brochet, Jerôme Lane, et al. Imgt®, the international immunogenetics information system®. Nucleic acids research, 37(Suppl 1):D1006–D1012, 2009.

[22] Vanessa Mhanna, Habib Bashour, Khang L ê Quy, Pierre Barennes, Puneet Rawat, Victor Greiff, and Encarnita Mariotti-Ferrandiz. Adaptive immune receptor repertoire analysis. Nature Reviews Methods Primers, 4(1):6, 2024.

[23] Ivana Mikocziova, Moriah Gidoni, Ida Lindeman, Ayelet Peres, Omri Snir, Gur Yaari, and Ludvig M Sollid. Polymorphisms in human immunoglobulin heavy chain variable genes and their upstream regions. Nucleic Acids Research, 48(10):5499–5510, 2020.

[24] Ivana Mikocziova, Ayelet Peres, Moriah Gidoni, Victor Greiff, Gur Yaari, and Ludvig M Sollid. Germline polymorphisms and alternative splicing of human immunoglobulin light chain genes. Iscience, 24(10), 2021.

[25] Supriya Munshaw and Thomas B. Kepler. Soda2: a hidden markov model approach for identification of immunoglobulin rearrangements. Bioinformatics, 26(7):867–872, February 2010.

[26] Kenneth Murphy and Casey Weaver. Janeway’s immunobiology. Garland science, 2016.

[27] Valerie H Odegard and David G Schatz. Targeting of somatic hypermutation. Nature Reviews Immunology, 6(8):573–583, 2006.

[28] Aviv Omer, Ayelet Peres, Oscar L Rodriguez, Corey T Watson, William Lees, Pazit Polak, Andrew M Collins, and Gur Yaari. T cell receptor beta germline variability is revealed by inference from repertoire data. Genome medicine, 14:1–19, 2022.

[29] Aviv Omer, Or Shemesh, Ayelet Peres, Pazit Polak, Adrian J Shepherd, Corey T Watson, Scott D Boyd, Andrew M Collins, William Lees, and Gur Yaari. Vdjbase: an adaptive immune receptor genotype and haplotype database. Nucleic acids research, 48(D1):D1051– D1056, 2020.

[30] Matt Pennell, Oscar L Rodriguez, Corey T Watson, and Victor Greiff. The evolutionary and functional significance of germline immunoglobulin gene variation. Trends in immunology, 44(1):7–21, 2023.

[31] Ayelet Peres, Moriah Gidoni, Pazit Polak, and Gur Yaari. Rabhit: R antibody haplotype inference tool. Bioinformatics, 35(22):4840–4842, 2019.

[32] Ayelet Peres, William D Lees, Oscar L Rodriguez, Noah Y Lee, Pazit Polak, Ronen Hope, Meirav Kedmi, Andrew M Collins, Mats Ohlin, Steven H Kleinstein, Corey T Watson, and Gur Yaari. IGHV allele similarity clustering improves genotype inference from adaptive immune receptor repertoire sequencing data. Nucleic Acids Research, page gkad603, 08 2023.

[33] Christelle Pommié, Séverine Levadoux, Robert Sabatier, Gérard Lefranc, and Marie-Paule Lefranc. IMGT standardized criteria for statistical analysis of immunoglobulin V-REGION amino acid properties. J. Mol. Recognit., 17(1):17–32, January 2004.

[34] Pradeepa Pushparaj, Andrea Nicoletto, Daniel J Sheward, Hrishikesh Das, Xaquin Castro Dopico, Laura Perez Vidakovics, Leo Hanke, Mark Chernyshev, Sanjana Narang, Sungyong Kim, et al. Immunoglobulin germline gene polymorphisms influence the function of sars-cov-2 neutralizing antibodies. Immunity, 56(1):193–206, 2023.

[35] Duncan K. Ralph and Frederick A. Matsen. Consistency of vdj rearrangement and substitution parameters enables accurate b cell receptor sequence annotation. PLOS Computational Biology, 12(1):e1004409, January 2016.

[36] Oscar L Rodriguez, Yana Safonova, Catherine A Silver, Kaitlyn Shields, William S Gibson, Justin T Kos, David Tieri, Hanzhong Ke, Katherine JL Jackson, Scott D Boyd, et al. Genetic variation in the immunoglobulin heavy chain locus shapes the human antibody repertoire. Nature communications, 14(1):4419, 2023.

[37] Modi Safra, Lael Werner, Ayelet Peres, Pazit Polak, Naomi Salamon, Michael Schvimer, Batia Weiss, Iris Barshack, Dror S Shouval, and Gur Yaari. A somatic hypermutation–based machine learning model stratifies individuals with crohn’s disease and controls. Genome Research, 33(1):71–79, 2023.

[38] Geir Kjetil Sandve and Victor Greiff. Access to ground truth at unconstrained size makes simulated data as indispensable as experimental data for bioinformatics methods development and benchmarking. Bioinformatics, 38(21):4994–4996, 2022.

[39] David G Schatz and Yanhong Ji. Recombination centres and the orchestration of v (d) j recombination. Nature reviews immunology, 11(4):251–263, 2011.

[40] Erand Smakaj, Lmar Babrak, Mats Ohlin, Mikhail Shugay, Bryan Briney, Deniz Tosoni, Christopher Galli, Vendi Grobelsek, Igor D’Angelo, Branden Olson, Sai Reddy, Victor Greiff, Johannes Trück, Susanna Marquez, William Lees, and Enkelejda Miho. Benchmarking immunoinformatic tools for the analysis of antibody repertoire sequences. Bioinformatics, 36(6):1731–1739, December 2019.

[41] Omri Snir, Luka Mesin, Moriah Gidoni, Knut EA Lundin, Gur Yaari, and Ludvig M Sollid. Analysis of celiac disease autoreactive gut plasma cells and their corresponding memory compartment in peripheral blood using high-throughput sequencing. The Journal of Immunology, 194(12):5703–5712, 2015.

[42] Joel NH Stern, Gur Yaari, Jason A Vander Heiden, George Church, William F Donahue, Rogier Q Hintzen, Anita J Huttner, Jon D Laman, Rashed M Nagra, Alyssa Nylander, et al. B cells populating the multiple sclerosis brain mature in the draining cervical lymph nodes. Science translational medicine, 6(248):248ra107–248ra107, 2014.

[43] Jason Anthony Vander Heiden, Susanna Marquez, Nishanth Marthandan, Syed Ahmad Chan Bukhari, Christian E Busse, Brian Corrie, Uri Hershberg, Steven H Kleinstein, Frederick A Matsen IV, Duncan K Ralph, et al. Airr community standardized representations for annotated immune repertoires. Frontiers in immunology, 9:2206, 2018.

[44] Gur Yaari and Steven H Kleinstein. Practical guidelines for b-cell receptor repertoire sequencing analysis. Genome medicine, 7:1–14, 2015.

[45] Gur Yaari, Mohamed Uduman, and Steven H Kleinstein. Quantifying selection in high-throughput immunoglobulin sequencing data sets. Nucleic acids research, 40(17):e134–e134, 2012.

[46] Gur Yaari, Jason A. Vander Heiden, Mohamed Uduman, Daniel Gadala-Maria, Namita Gupta, Joel N. H. Stern, Kevin C. O’Connor, David A. Hafler, Uri Laserson, Francois Vigneault, and Steven H. Kleinstein. Models of somatic hypermutation targeting and substitution based on synonymous mutations from high-throughput immunoglobulin sequencing data. Frontiers in Immunology, 4, 2013.

[47] Christina Yacoob, Marie Pancera, Vladimir Vigdorovich, Brian G Oliver, Jolene A Glenn, Junli Feng, D Noah Sather, Andrew T McGuire, and Leonidas Stamatatos. Differences in allelic frequency and cdrh3 region limit the engagement of hiv env immunogens by putative vrc01 neutralizing antibody precursors. Cell reports, 17(6):1560–1570, 2016.

[48] Jian Ye, Ning Ma, Thomas L. Madden, and James M. Ostell. Igblast: an immunoglobulin variable domain sequence analysis tool. Nucleic Acids Research, 41(W1):W34–W40, May 2013.

[49] Yan Zhu, Haipei Tang, Wenxi Xie, Sen Chen, Huikun Zeng, Chunhong Lan, Junjie Guan, Cuiyu Ma, Xiujia Yang, Qilong Wang, et al. The multilevel extensive diversity across the cynomolgus macaque captured by ultra-deep adaptive immune receptor repertoire sequencing. Science Advances, 10(4):eadj5640, 2024.

