## supplementary information for "An unbiased comparison of immunoglobulin sequence aligners"

June 9, 2024

#### 1 Generating Non-Polymorphic Regions

To generate the nontemplated and palindromic (NP) regions, GenAIRR uses a Markov process derived from empirical data. The process is tailored individually for the first and second NP regions to accurately represent the unique characteristics of each segment. Mathematically, the Markov chain for an NP region operates as follows: given the current nucleotide $X_t$ at position $t$ in the NP region, the chain provides a probability distribution over the four possible nucleotides for the next position $X_{t+1}$. This probability distribution is conditional on the current nucleotide and its position within the NP region, expressed as:

$$P(X_{t+1} = x \mid X_t = x_t, t) = f(x_t, t)$$

where $f(x_t, t)$ denotes the function that determines the probability of each nucleotide based on the current nucleotide $x_t$ and the position $t$.

#### 2 Analysis of Minimal D Allele Length for Heavy Chain Classification

This analysis focuses on identifying the minimal nucleotide sequence length required to uniquely identify immunoglobulin heavy chain D alleles. By analyzing mutually exclusive substrings (MES) within each allele, we found that an MES of at least 11 nucleotides is required for unique identification of all alleles. Figure 1 (Panel A) shows that any substring shorter than this may not be unique, potentially leading to ambiguous classification.

Aligners face challenges in reliably identifying these alleles due to somatic hypermutation and sequencing inaccuracies. However, as shown in Figure 1 (Panel B), identifying around 8 nucleotides allows for accurate classification of approximately 90% of alleles, based on a theoretical perfect model. Below 8 nucleotides, classification accuracy drops sharply, indicating a trade-off between MES length and coverage.

To address the challenges of short D alleles post-recombination, GenAIRR introduces a hyperparameter threshold of 5 bases, covering roughly 85% of D alleles (see Supplementary Figure 1). This threshold balances computational efficiency with biological realism, enabling more reliable classification among the majority of D alleles.

Figure 1 (Panel C) provides an ECDF of 6M generated sequence D allele length, demonstrating the distribution of MES lengths across different alleles. This data emphasizes the importance of advanced alignment tools capable of accurately classifying these alleles despite varying MES lengths.

The analysis and simulations presented here highlight the need for careful consideration of D allele lengths and trimming effects to enhance the accuracy of sequence classification in practical scenarios.

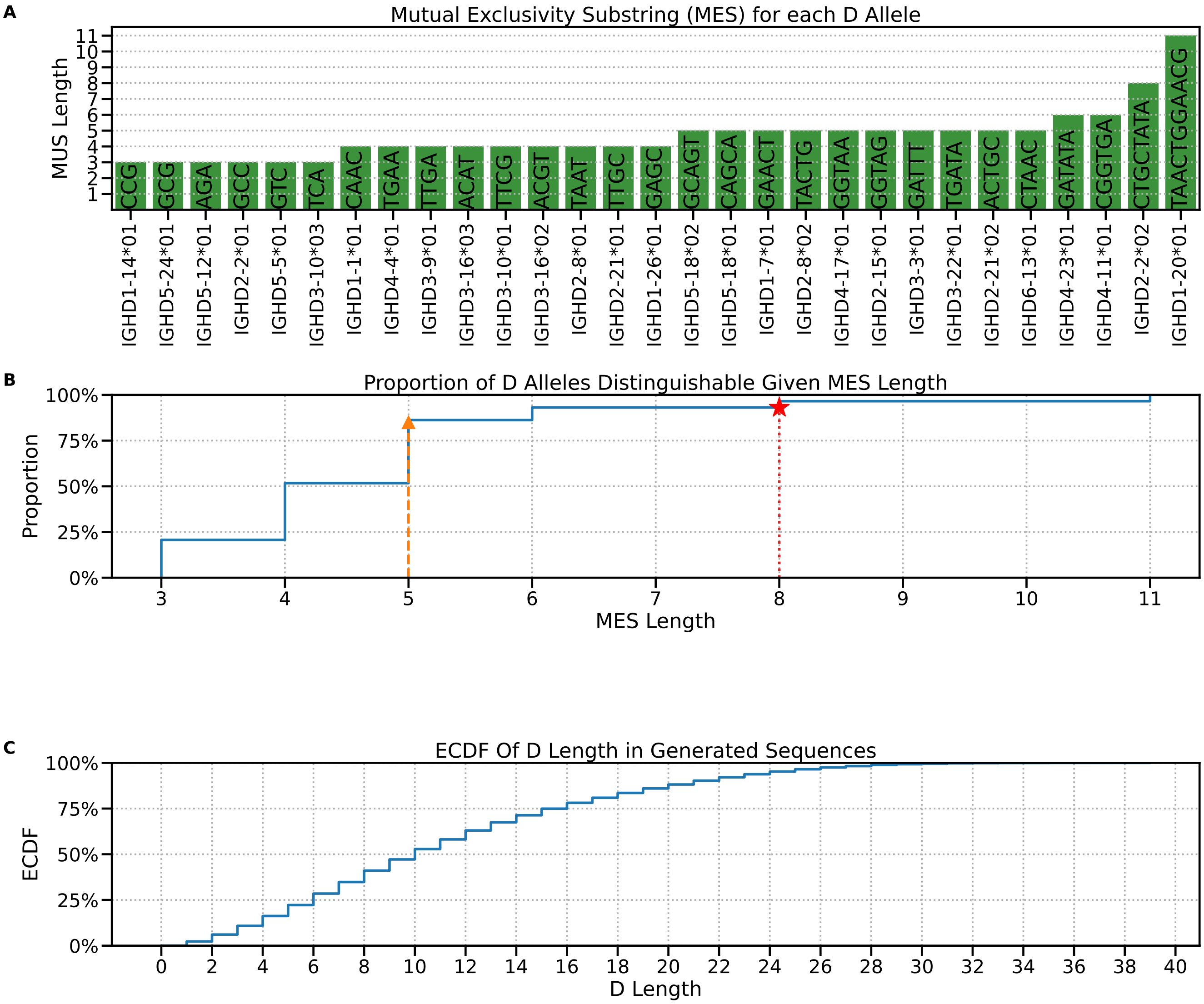


Figure 1: Panel A - Bar graph illustrating the shortest mutually exclusive substrings (MES) lengths for a set of immunoglobulin heavy-chain D alleles. Each green bar represents the length of the MES required to uniquely identify each corresponding D allele, indicated on the x-axis, with the actual MES sequence displayed within the bar. The tallest bar reaches a length of 11, demonstrating the minimum length necessary to differentiate all alleles in the set uniquely. Panel B - illustrates coverage thresholds with a red line at 8 nucleotides and an orange line at 5 nucleotides, which correspond to approximately 90% and 83% of the reference set being uniquely classified, respectively, indicated by red stars and orange triangles on the graph. Panel C- The ECDF of 6M generated sequence D allele length

#### 2.1 Ambiguity Resolution and Ground Truth Alignment in Sequence Simulation

During the sequence generation process in GenAIRR, potential ambiguities that may arise due to the trimming or corruption of allele ends (5' or 3') are meticulously tracked and resolved. This involves the careful alignment and adjustment of the ground truth to accurately reflect the impact of these events. One common type of ambiguity occurs when critical distinguishing regions of an allele are removed—either due to deliberate trimming or unintended corruption. Such removal can render it impossible to distinguish between two or more alleles, as the unique identifiers that differentiate them are lost. To address this, a simulation schema is employed by GenAIRR that infers a map object post-trimming. This object identifies which alleles become indistinguishable under specific trimming scenarios, allowing the simulation to extend the number of correct calls for that sequence in the ground truth.

Additionally, corrections are applied by GenAIRR to the start and end positions of each allele in the ground truth. Specifically, after the recombination process and the simulated insertion of NP regions to bridge the junctions, the simulation schema assesses whether the generated NP region has inadvertently recreated an exact part of the allele sequence that was removed during the trimming process. If such a recreation occurs, the start or end positions of the alleles in the ground truth are updated to accurately reflect the new sequence configuration. This meticulous approach ensures that the integrity of the ground truth is maintained, enabling more precise and reliable simulation outcomes.

#### 2.2 Simulation of Corruption Events in GenAIRR

A distinctive feature of the GenAIRR simulation framework is its ability to simulate "corruption" events, which are designed to emulate some of the common errors encountered during sequencing. In GenAIRR, we simulate three main types of corruption events affecting the 5' end of the V gene: the Add event, the Remove event, and the Remove then Add event. Each type of event, as indicated by its name, involves either the addition or removal of sequence elements, or a combination of both.

The occurrence of these corruption events, along with the type of event that occurs, can be finely controlled using flexible parameters defined in the simulation arguments file. Both the removal and addition events are governed by pre-calculated distributions based on empirical data, which determine the likelihood of observing a specific length of removal or addition. GenAIRR allows users to adjust these distributions through three coefficient parameters, enabling the customization of the extent of sequence alterations, such as shorter removals or additions, as needed for specific simulation scenarios. Furthermore, under the Add event, GenAIRR supports four subtypes of additions:

- **Random Addition:** After determining the length of the addition from the distribution, a random DNA sequence of that length is generated.
- **Single Stream Addition:** This involves choosing one nucleotide, including 'N', and repeating it to create a string of the sampled length.
- **Random Allele Section:** This event entails randomly selecting an allele from the reference file and then choosing a random section of that allele equal to the length determined for the addition.
- **Start Duplication:** Based on the sampled length, this event duplicates a corresponding segment from the start of the current sequence to use as the addition.

Regardless of the type of addition event chosen, all resultant addition strings are concatenated to the beginning of the sequence. For the purposes of this study, the simulation parameters were specifically set to employ the random sequence addition event exclusively, to maintain consistency and control within the experimental framework. This approach allows us to assess the impact of sequencing errors on the alignment and interpretation of immunoglobulin sequences in a controlled manner, providing insights into the robustness of different sequence aligners in the face of simulated sequencing errors.

### 3 GenAIRR Productivity Testing Scheme

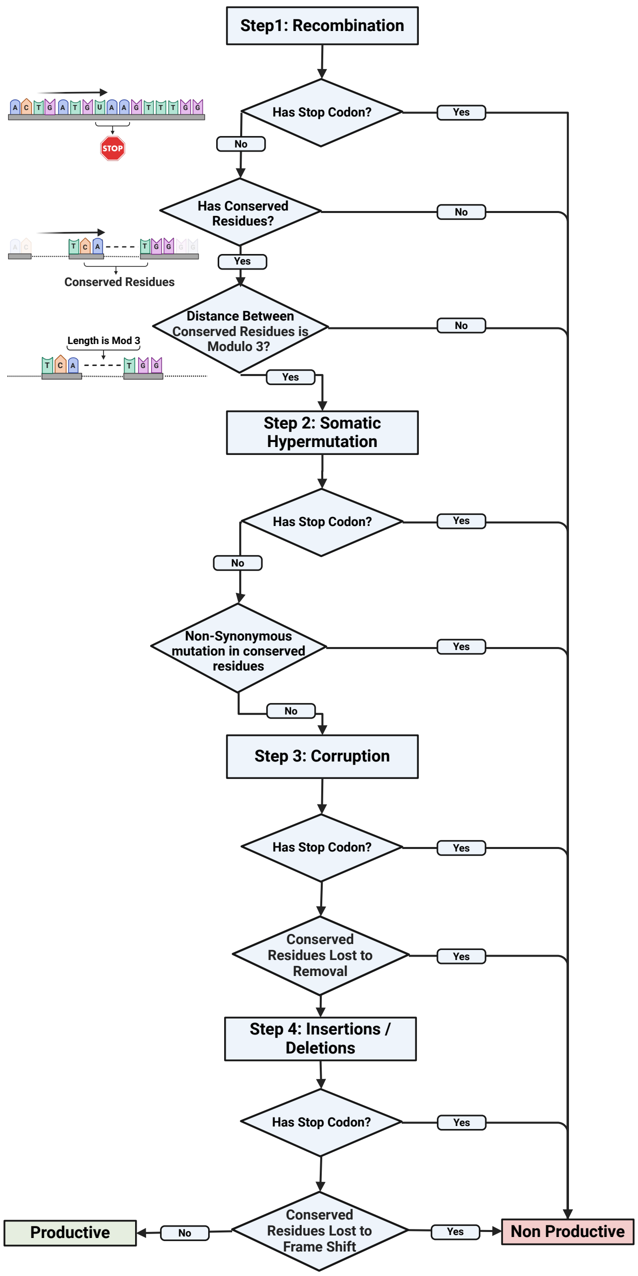


Figure 2: Schematic diagram outlining the productivity assessment in GenAIRR generated sequences. The flowchart details four stages at which GenAIRR verifies productivity of a sequence: 1) Recombination, where sequences are evaluated for stop codons and the presence of conserved residues; 2) Somatic hypermutation, assessing the impact of non-synonymous mutations on conserved residues and their spacing;3)After applying corruption to the 5' end of the ✓gene; 4) Insertions/Deletions, determining sequence productivity based on frame shifts and the loss of conserved residues. Each step incorporates decision points that define whether a sequence is considered productive or non-productive.

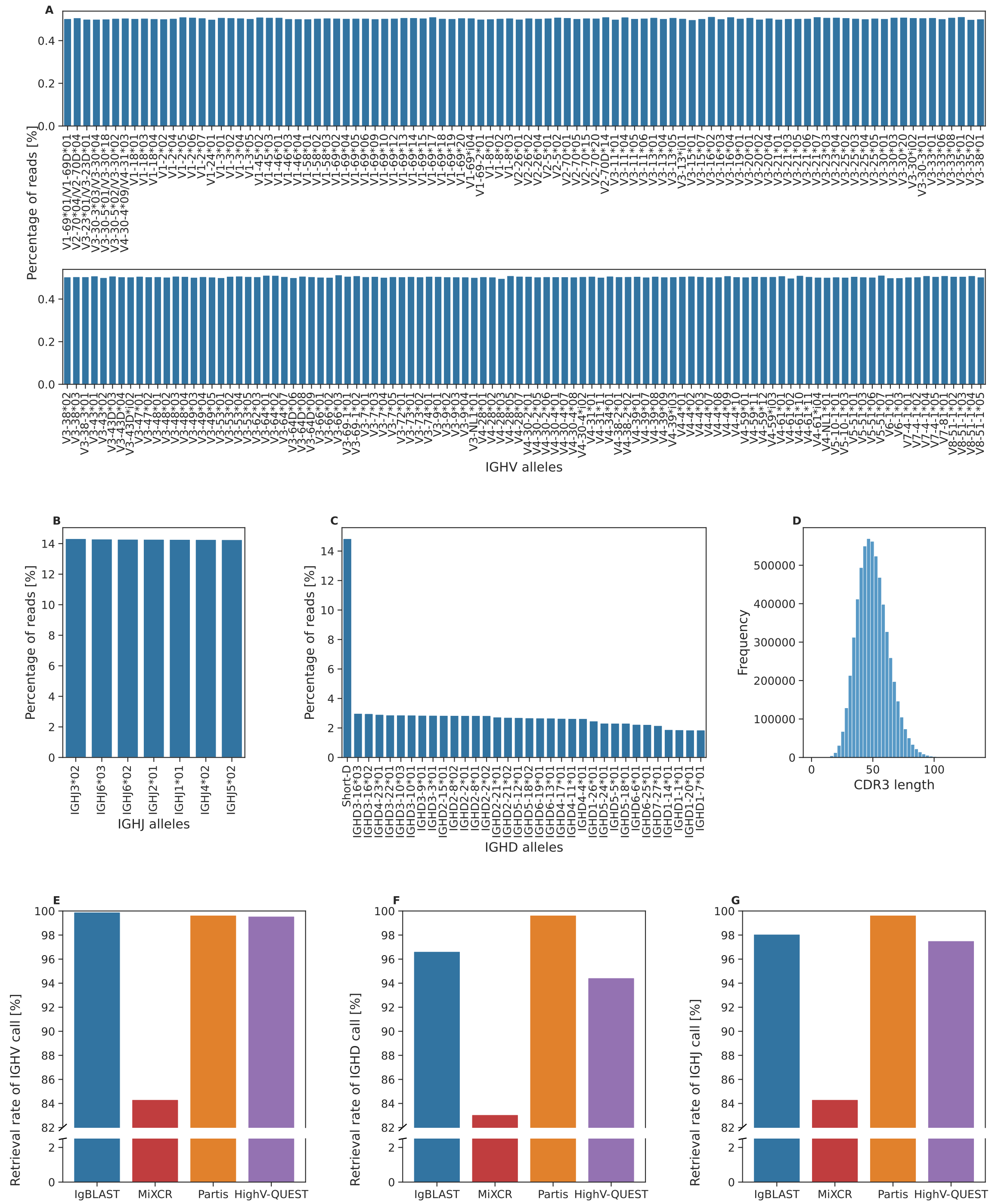


Figure 3: Overview of the simulated nonproductive dataset
(A-C) Distribution of the V, D, and J allele usage in the nonproductive dataset. Each column represents a different allele, and the y-axis indicates their relative usage percentage in the dataset. (D) CDR3 length distribution. The x-axis shows the CDR3 lengths, while the y-axis indicates their frequency. (E-G) Aligner's allele assignment coverage. Each column represents a different aligner, and the y-axis shows the percentage of sequences for which the aligner returned an allele assignment. The colors correspond to the different aligners: blue for IgBLAST, red for MiXCR, orange for Partis, and purple for HighV-QUEST.

Table 1: Comparison of IG Sequence Simulators

| **Feature/Parameter** | **GenAIRR** | **AIRRSHIP** | **IGoR** | **OLGA** | **immuneSIM** | **IMPlAntS** | **partis** | **Shmulate** |
|---|---|---|---|---|---|---|---|---|
| **Ambiguity Resolution** | ✓ | ✗ | ✗ | ✗ | ✗ | ✗ | ✗ | ✗ |
| **Region Reconstruction Validation** | ✓ | ✗ | ✗ | ✗ | ✗ | ✗ | ✗ | ✗ |
| **Dynamic Start/End Modification** | ✓ | ✗ | ✗ | ✗ | ✗ | ✗ | ✓ | ✗ |
| **Sequencing Error Emulation** | ✓ | ✗ | ✗ | ✗ | ✗ | ✓ | ✗ | ✗ |
| **SHM Simulation** | ✓ | ✓ | ✓ | ✓ | ✓ | ✓ | ✓ | ✓ |
| **Diverse Mutation Model** | ✓ | ✓ | ✗ | ✗ | ✓ | ✓ | ✓ | ✓ |
| **Tunable Mutation Distributions** | ✓ | ✓ | ✗ | ✗ | ✓ | ✓ | ✓ | ✓ |
| **Simulation Modularity** | ✓ | ✓ | ✗ | ✗ | ✓ | ✓ | ✗ | ? |
| **"Ns" Simulation** | ✓ | ✓ | ✓ | ✓ | ✗ | ✓ | ✗ | ✗ |
| **Recorded Mutation Events** | ✓ | ✗ | ✗ | ✗ | ✗ | ✗ | ✗ | ✗ |
| **Scalability** | ✓ | ✓ | ✓ | ✓ | ✓ | ? | ✓ | ✓ |
| **Custom Reference** | ✓ | ✗ | ✓ | ✗ | ✗ | ✓ | ✓ | ✓ |
| **Indel Simulation** | ✓ | ✗ | ✓ | ✓ | ✓ | ✓ | ✓ | ✗ |
| **Support for Different Chains** | ✓ | ✗ | ✓ | ✓ | ✓ | ✗ | ✓ | ✓ |
| **Output Format Options** | ✓ | ✓ | ✓ | ✓ | ✓ | ✓ | ✗ | ✗ |
| **Documentation** | ✓ | ✓ | ✓ | ✓ | ✓ | ✗ | ✓ | ✓ |
| **Under Support** | ✓ | ✓ | ✓ | ✓ | ? | ? | ✓ | ✓ |
| **Productivity Testing** | ✓ | ✓ | ✗ | ✗ | ✗ | ✓ | ✓ | ✗ |
| **Open Source** | ✓ | ✓ | ✓ | ✓ | ✓ | ✓ | ✓ | ✓ |
| **Available Code** | ✓ | ✓ | ✓ | ✓ | ✓ | ✓ | ✓ | ✓ |
| **Platform availability** | Standalone | Standalone | Standalone | Standalone | Standalone | Standalone | Standalone | Standalone |
| **Release Year** | 2023 | 2023 | 2018 | 2019 | 2020 | 2021 | 2016 | 2017 |
| **Last updated** | 2024 | 2023 | 2020 | 2021 | 2021 | 2023 | 2024 | 2024 |

Table 2: Extended Comparison of Different IG Sequence Aligners

| **Criteria** | **IgBlast** | **iHMMune-align** | **BRILIA** | **SoDA2** | **IgSCUEAL** | **IMSEQ** | **Joinsolver** | **MiXCR** | **partis** | **VDJFasta** | **VDJsolver** | **HighV-QUEST** | **ImReP** | **IgMAT** | **abstar** | **AbAlign** |
|---|---|---|---|---|---|---|---|---|---|---|---|---|---|---|---|---|
| Release Year | 2013 | 2007 | 2017 | 2010 | 2015 | 2015 | 2004 | 2015 | 2016 | 2009 | 2006 | 2008 | 2020 | 2023 | 2016 | 2023 |
| Last Updated | 2023 | 2009 | 2018 | - | 2016 | 2016 | - | 2024 | 2024 | 2015 | 2006 | 2022 | 2021 | 2023 | 2023 | 2024 |
| Available Code | ✓ | ✓ | ✓ | Site not available | ✓ | ✓ | ✗ | ✓ | ✓ | ✓ | ✓ | ✗ | ✓ | ✓ | ✓ | ✓ |
| Platform availability | Online/ Stand-alone | Online/ Stand-alone | Stand-alone | Stand-alone | Stand-alone | Stand-alone | Stand-alone | Stand-alone | Stand-alone | Stand-alone | Online/ Stand-alone | Online | Stand-alone | Stand-alone | Stand-alone | Stand-alone |
| Underlining Method | Blast | HMM | MSA | HMM | MSA | SCF matching and alignment | CDR3 alignment | MSA | HMM | HMM | MSA | MSA | CDR3 alignment | HMM and MSA | Blast | MSA |
| Gene / Alleles | Alleles | Alleles | Genes | Alleles | Alleles | Genes | Alleles | Alleles | Alleles | Alleles | Alleles | Alleles | Alleles | Regions | Alleles | Genes |
| Custom reference | ✓ | ✓ | ✓ | ✓ | ✓ | ✓ | ✗ | ✓ | ✓ | ✓ | ✗ | ✗ | ✓ | ✓ | ✓ | ? |
| Programming Language | C++ | Java | Matlab | - | Python and JS | C++ | - | Java | Python | Perl | - | - | Python | Python | Python | C++ |

### 4 GenAIRR Parameters used in Manuscript Dataset

Table 3: Parameters used in GenAirr to generate each dataset.

| **Parameter** | **Productive Dataset** | **Non-Productive Dataset** |
|---|---|---|
| **Min Mutation Rate** | 0.003 | 0.003 |
| **Max Mutation Rate** | 0.25 | 0.25 |
| **Simulate Indels** | 0 | 0.2 |
| **Max Indels** | 5 | 5 |
| **Deletion Probability** | 0.5 | 0.5 |
| **Insertion Probability** | 0.5 | 0.5 |
| **N Ratio** | 0 | 0.02 |
| **N Probability** | 0 | 0.2 |
| **Max Sequence Length** | 512 | 512 |
| **Mutation Model** | Uniform | Uniform |
| **Custom Mutation Model Path** | None | None |
| **Nucleotide Add Coefficient** | 210 | 210 |
| **Nucleotide Remove Coefficient** | 310 | 310 |
| **Nucleotide Add After Remove Coefficient** | 50 | 50 |
| **Random Sequence Add Probability** | 1 | 1 |
| **Single Base Stream Probability** | 0 | 0 |
| **Duplicate Leading Probability** | 0 | 0 |
| **Random Allele Probability** | 0 | 0 |
| **Corrupt Probability** | 0 | 0.7 |
| **Corrupt Events Probability** | [0.5, 0.5, 0] | [0.5, 0.5, 0] |
| **Short D Length** | 5 | 5 |
| **Kappa Lambda Ratio** | 0.5 | 0.5 |
| **Save Mutations Record** | Yes | Yes |
| **Save Ns Record** | Yes | Yes |
| **Save Corruption Record** | No | No |
| **Productive** | Yes | No |

### 5 Output Format of GenAIRR Datasets

Table 4: Example of Output Format File of GenAIRR

| Field Name | Sequence 1 | Sequence 2 | Sequence 3 | Sequence 4 |
|---|---|---|---|---|
| **sequence** | GATTCACCTT... | GAGGTGCAGC... | GGTCCGCCAN... | CCCCTCCCGC... |
| **v_sequence_start** | 0 | 0 | 0 | 0 |
| **v_sequence_end** | 220 | 293 | 185 | 114 |
| **d_sequence_start** | 220 | 293 | 202 | 118 |
| **d_sequence_end** | 229 | 302 | 203 | 122 |
| **j_sequence_start** | 246 | 312 | 258 | 183 |
| **j_sequence_end** | 295 | 372 | 258 | 183 |
| **v_germline_start** | 76 | 0 | 110 | 185 |
| **v_germline_end** | 296 | 296 | 295 | 299 |
| **d_germline_start** | 0 | 0 | 10 | 8 |
| **d_germline_end** | 9 | 9 | 11 | 12 |
| **j_germline_start** | 2 | 3 | 1 | 0 |
| **j_germline_end** | 51 | 63 | 50 | 50 |
| **junction_sequence_start** | 209 | 282 | 175 | 103 |
| **junction_sequence_end** | 264 | 341 | 227 | 152 |
| **v_call** | IGHVF10-G49*03 | IGHVF4-G14*03 | IGHVF4-G14*01 | IGHVF4-G1*01 |
| **d_call** | IGHD6-6*01 | IGHD6-6*01, IGHD6-6*02 | Short-D | Short-D |
| **j_call** | IGHJ5*02 | IGHJ6*02 | IGHJ3*02 | IGHJ3*02 |
| **mutation_rate** | 0.064 | 0.086 | 0.058 | 0.158 |
| **v_trim_5** | 0 | 0 | 0 | 0 |
| **v_trim_3** | 0 | 0 | 0 | 0 |
| **d_trim_5** | 0 | 1 | 10 | 9 |
| **d_trim_3** | 9 | 10 | 5 | 6 |
| **j_trim_5** | 2 | 3 | 1 | 0 |
| **j_trim_3** | 0 | 0 | 0 | 0 |
| **corruption_event** | remove | no-corruption | remove | remove |
| **corruption_add_amount** | 0 | 0 | 0 | 0 |
| **corruption_remove_amount** | 76 | 0 | 110 | 185 |
| **indels** | {} | {14: D > C, 211: D > G, 243: D > T} | {} | {} |
| **productive** | False | False | False | False |
| **stop_codon** | True | True | True | True |
| **vj_in_frame** | False | False | False | False |
| **note** | | | | |
